# Effects of partial, combined and total replacement of sodium chloride in beef sausage on microbial load, sensory acceptability and physical properties

**DOI:** 10.1101/506071

**Authors:** Olusegun D. Oshibanjo, Odion O. Ikhimiukor

## Abstract

This study evaluates the effects of partial, combined or total replacement of NaCl on microbial load, sensory evaluation and physical properties of beef sausage. Beef sausage was prepared (g/100g: beef 65.0, corn flour 10.0, oil 10.0, soya bean, wet and dry spice 15.0). Sodium chloride (SC) was replaced with potassium chloride (PC), potassium lactate (PL) or calcium ascorbate (CA) at 25%, 50% and 100%, and then stored for 15 days in a factorial arrangement in complete randomized design. Sausages were subjected to microbial load, sensory evaluation and pH, cooking yield and cooking loss using standard procedure. Data were analysed using descriptive statistic and ANOVA at α0.05. The microbial load was generally lower. The pH increased with storage time. The cooking yield was significantly higher in salt combination of 50% SC and CA each at storage day 5 with least cooking loss at same storage day. The most preferred sausage colour was obtained with a salt combination of 50% SC and CA each with least colour in a salt combination of 50% SC, 25% PL and). In tenderness, a salt combination of 50% of both SC and CA was significantly tender with least tenderness in 100% PL total replacement. The panelists rated salt combination of 50% SC and CA as the most overall accepted salt combination than others salt combination. Sodium chloride replacement with Calcium Ascorbate at 50% enhanced the most preferred sausage for overall acceptability and aerobic bacteria count lower than the international standard limit.

**Summary statement:** High intake of sodium chloride had been seen as one of the main contributing factors in the development of specific, non-transmittable diseases, such as, high blood pressure and related secondary such as cardiovascular disease. The use of other derivative of salt is therefore needed. This study evaluates the effects of partial, combined or total replacement of NaCl on microbial load, sensory evaluation and physical properties of beef sausage.

## Introduction

A link between increased salt consumption in sodium chloride (NaCl) and increased blood pressure has been verified in several studies (Sarmugam et al., *2014*). The Global Strategy on Diet, Physical Activity and Health passed by the World Health Organization (WHO, 2004). identified increased salt consumption as one of the main contributing factors in the development of specific, non-transmittable diseases, such as, high blood pressure and related secondary such as cardiovascular disease. Generally, most of the sodium in the human body comes from salt added during food processing. Many types of processed foods contribute to the high intake of sodium. Highly and healthier products will be rated higher in taste, only as long as the consumers do not have to sacrifice palatability. Furthermore, in many applications, the total exclusion of salt is not possible.

Therefore, adequate salt substitutes must feature essential functionalities in taste, preservation, texture and the color of end products. Importantly, the addition of salts in food has for a long time been used to inhibit the activity of microorganisms, particularly in retarding microbial food spoilage and/or food poisoning processes (Rawat, 2015)

Due to the importance of salts in food processing, the replacement of sodium salts with other less harmful salts is imperative. Replacement of Sodium chloride with potassium chloride has been reported and it’s the most commonly used salt to date (Weiss et al., 2010). However, potassium chloride has a slightly bitter taste, however, to prevent the product from having unacceptable sensory properties, masking substances are added as additives to the product. Similarly, flavour enhancers may also be added to the meat product. Flavour enhancers themselves do not have a salty taste, but may in combination with salt increase the saltiness of the product. For example, carboxymethylcellulose and carrageenan in combination with sodium citrate have been shown to enhance saltiness in frankfurters (Weiss et al., 2010 and Ruusunen et al., 2003a).

This study therefore seeks to evaluate the effect of partial, combined and total replacement of sodium chloride in beef sausage on microbial load, sensory acceptability and physical properties.

## Materials and methods

### Location of the study

The experiment was undertaken at the Animal Product and Processing (Meat Science) Laboratory of the Department of Animal Science, Faculty of Agriculture, University of Ibadan, Ibadan.

### Beef sausage formulation

Beef sausage was prepared using Table 1.

**Table 1:**
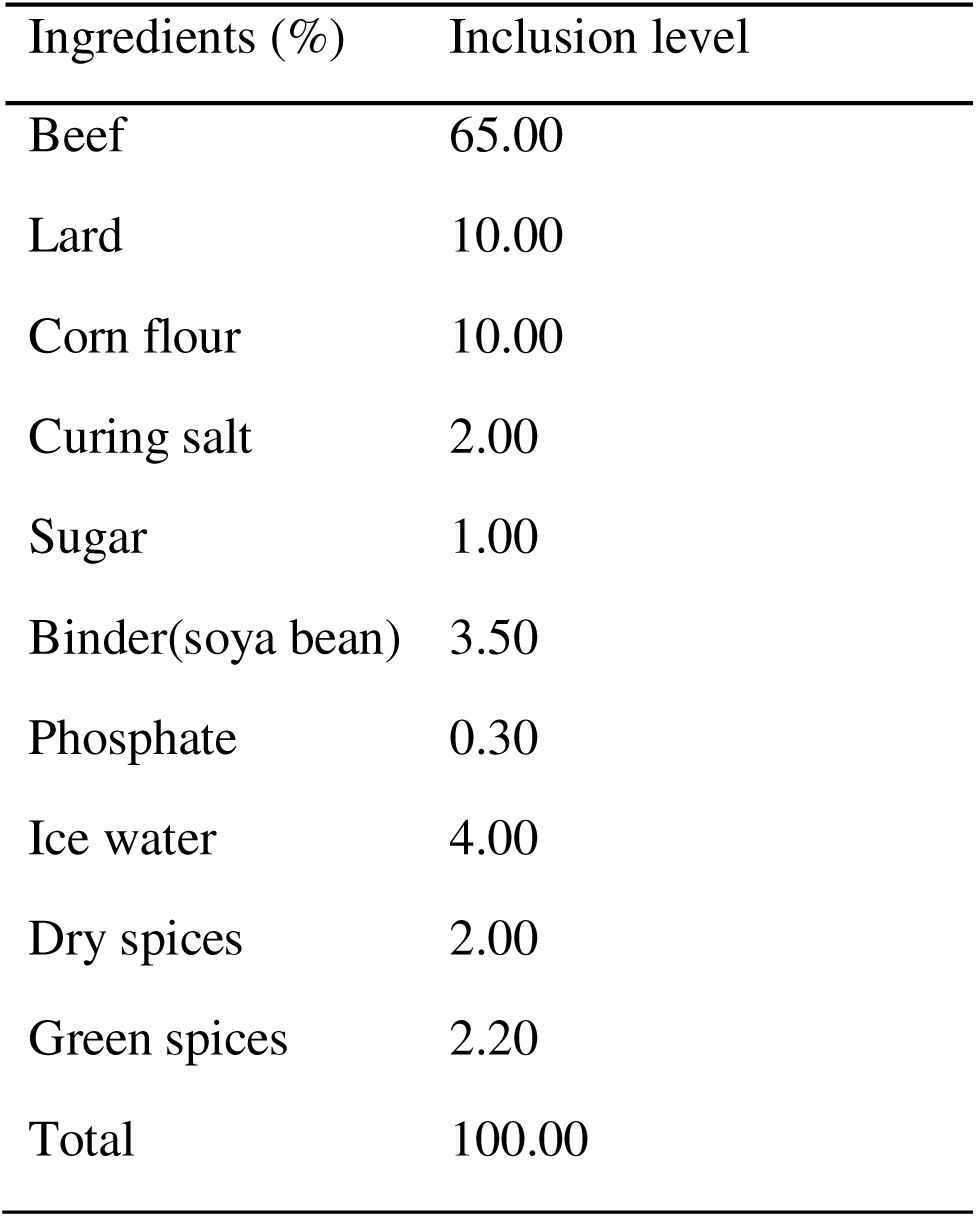
Beef sausage formulation

### Salt combination inclusion

Sodium chloride (SC) was replaced with potassium chloride (PC), potassium lactate (PL) or calcium ascorbate (CA) at 25, 50 and 100% stored at −4°C for 15 days as showed in Table 2.

**Table 2:**
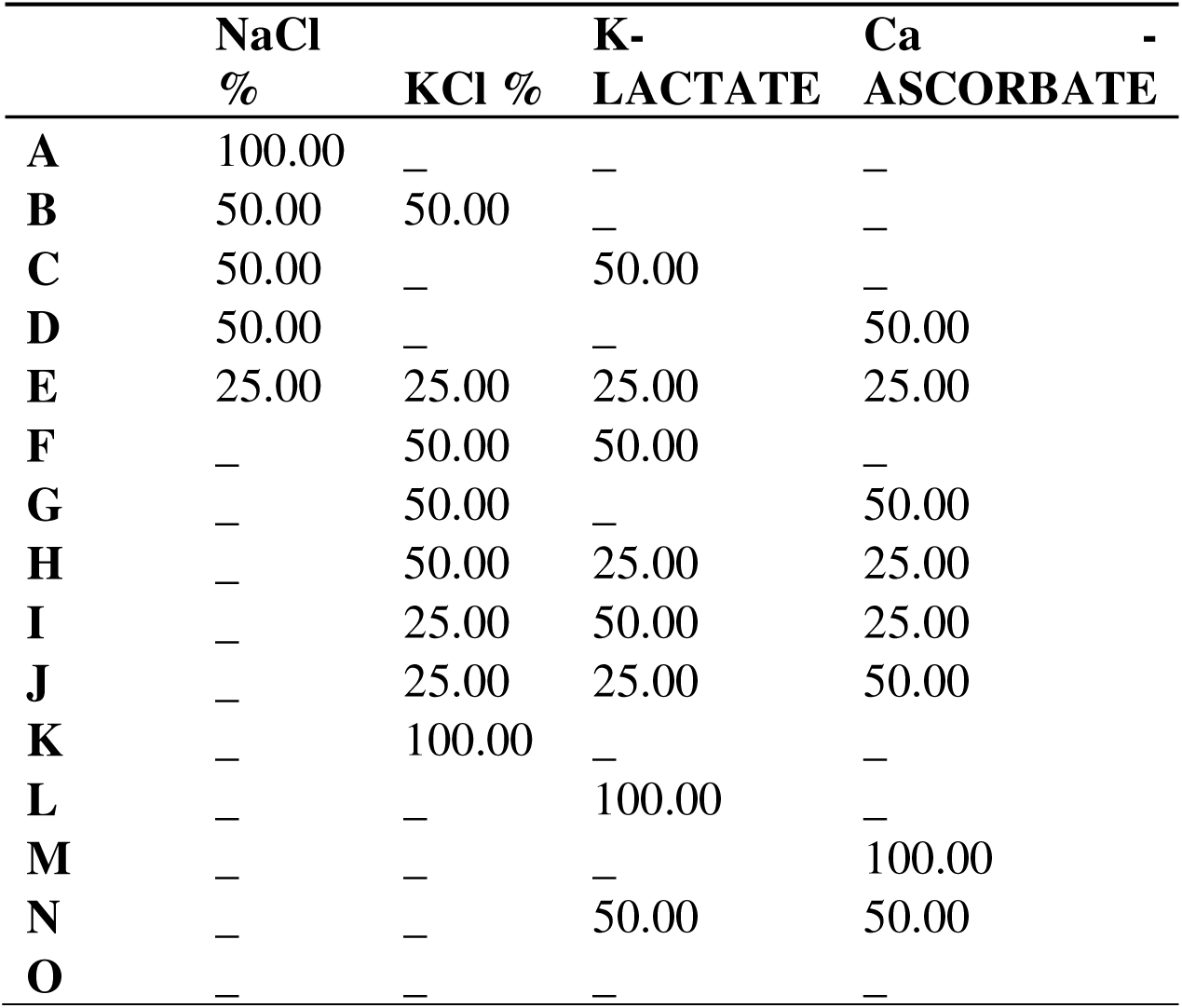
salt combination inclusion levels

### Microbial load evaluation

Microbial count was carried out using the pour plate method (Harrigan and McCanee 1976). A 1 ml dilutions of 10^−2^,10^−4^ and 10^−6^ were pipetted into molten agar at 45 °C, swirled and then introduced into sterile Petri dishes and incubated for 24 hours.

### Sensory evaluation

The nine-point hedonic scale was used by twenty panelists who were trained individuals aged between 20 and 40 years were used to determined two replicate of the prepared sausage to assess colour (1-4 dark, 5-intermediate, 6-9 light), tenderness (1-4 tough, 5-intermediate, 6-9 tender), juiciness (1-4 dry, 5-intermediate, 6-9 juicy), hotness (1-4 less, 5-intermediate, 6-9 high) and overall acceptability, OA (1-4 low, 5-intermediate, 6-9 high) Mahendrakar et al., 1988).

### Experimental design

Factorial arrangement in complete randomized design.

### Statistical Analysis

Data were subjected to analysis of variance using Statistical Analysis System SAS, (2002). Means were separated using Duncan’s Multiple Range Test option of the same software.

## Results

Table 3 shows the effects of different salts on the microbial count. There were no significant difference (P<0.05) in all the treatments across the storage days.

**Table 3:**
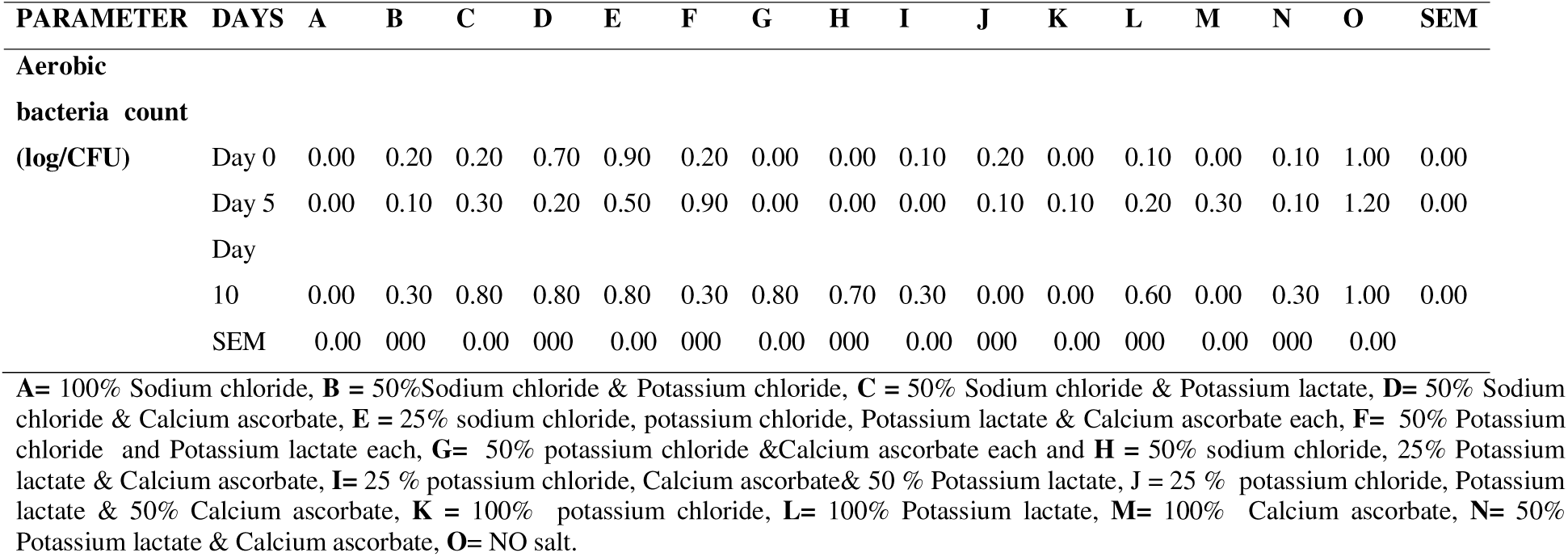
Interaction effect of different salt and storage days on microbial count of beef sausage (Log10CFU/ml)

The interaction effect of salt and storage days on pH value of beef sausage was showed on Table 4. There were significant difference (P>0.05) in all the treatments across the storage days. Treatment B and O containing 50%Sodium chloride and Potassium chloride and no salt respectively had the highest pH values at day 15 compared to treatment. It was generally observed that as the storage days increased there was increase in pH values obtained. Table 5 shows the interaction of salt and storage days on cooking yield of beef sausage. The cooking yield was significantly higher (P>0.05) in Treatment **D (**50% Sodium chloride and Calcium ascorbate) at day 5with least cooking yield in Treatment **L (**100% Potassium lactate) at day 10. While the cooking loss was significant higher in Treatment **L (**100% Potassium lactate) at day 10 with least cooking loss obtained in Treatment **D (**50% Sodium chloride and Calcium ascorbate) at day 5.

**Table 4:**
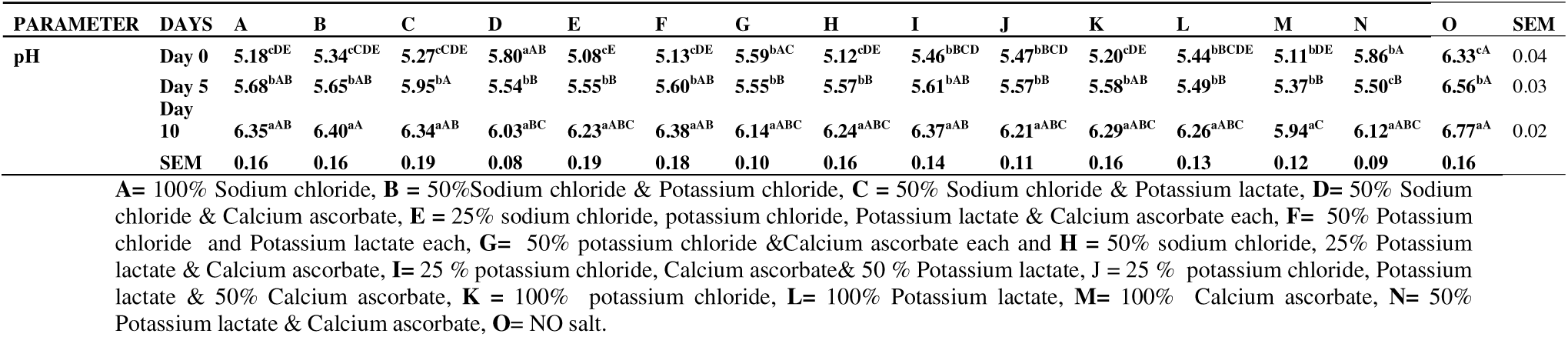
Interaction effect of different salt and storage days on pH of beef sausage

**Table 5:**
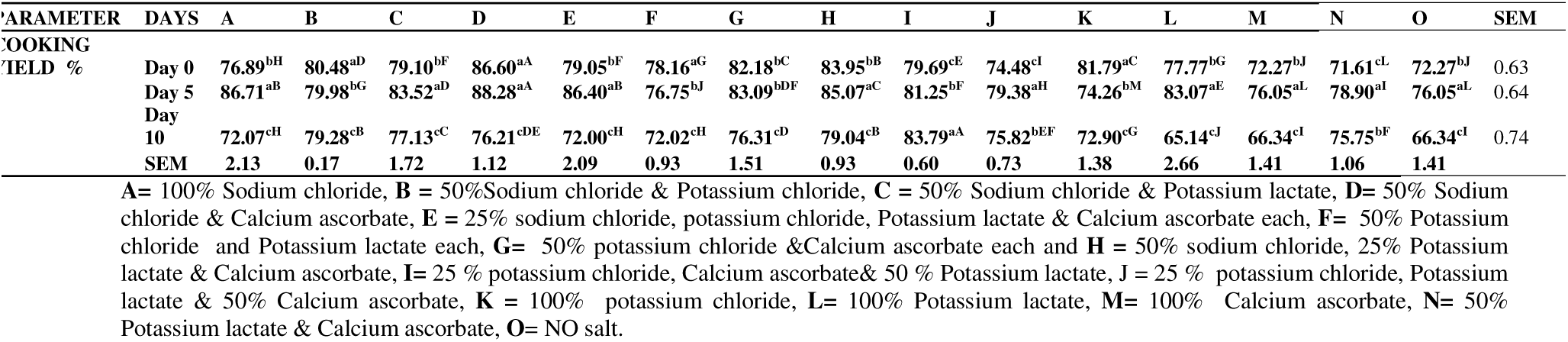
Interaction effect of different salt and storage days on cooking yield of beef sausage

Effect of different salts on sensory evaluation are shown on Figure 1 to 5. The most preferred sausage colour was obtained with a salt combination of 50% SC and CA each (5.60±0.2) with least colour in a salt combination of 50% SC, 25% PL and CA (2.60±0.1). For tenderness, a salt combination of 50% SC and CA each was significantly tender (6.60±0.3) with least tenderness in 100% PL total replacement. Salt combination of 25% SC, PC, PL and CA each was most juicy (6.30±0.3) while 100% PC total replacement was dry (3.80±0.1). The panelists rated salt combination of 50% SC and CA each as the most overall accepted salt combination (6.8±0.3) with least overall acceptability recorded for a salt combination of 50% PL and CA (3.20±0.1).

**Figure 1:**
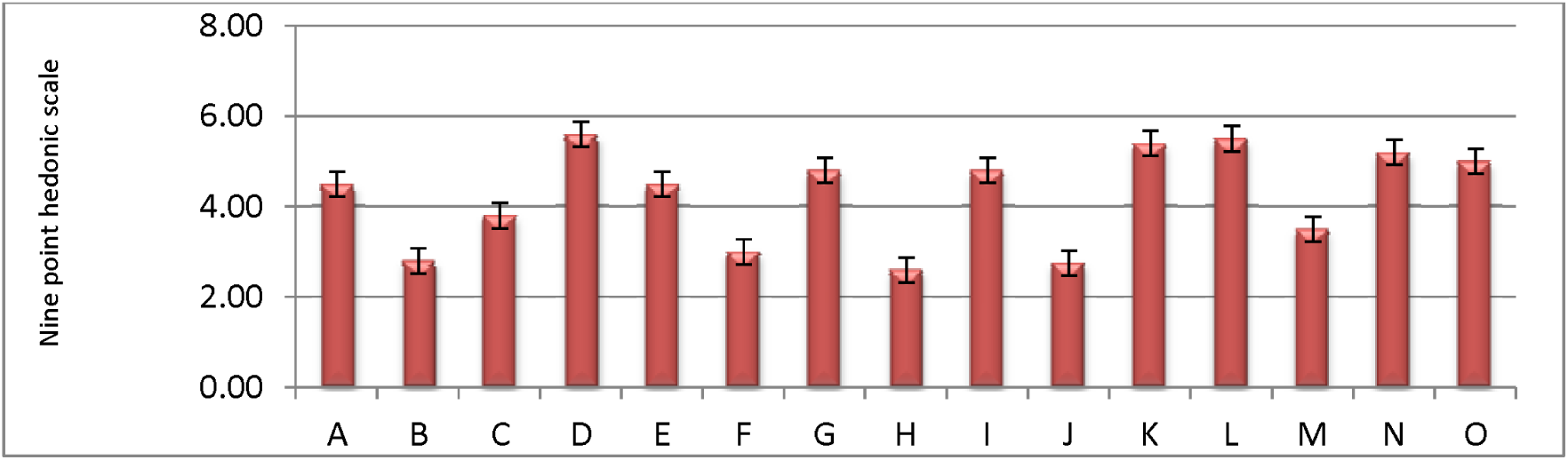
Effect of different salts on sausage colour. **A=** 100% Sodium chloride, **B =** 50%Sodium chloride & Potassium chloride, **C =** 50% Sodium chloride & Potassium lactate, **D=** 50% Sodium chloride & Calcium ascorbate, **E =** 25% sodium chloride, potassium chloride, Potassium lactate & Calcium ascorbate each, **F=** 50% Potassium chloride and Potassium lactate each, **G=** 50% potassium chloride &Calcium ascorbate each and **H =** 50% sodium chloride, 25% Potassium lactate & Calcium ascorbate, **I=** 25 % potassium chloride, Calcium ascorbate& 50 % Potassium lactate, J = 25 % potassium chloride, Potassium lactate & 50% Calcium ascorbate, **K =** 100% potassium chloride, **L=** 100% Potassium lactate, **M=** 100% Calcium ascorbate, **N=** 50% Potassium lactate & Calcium ascorbate, **O**= NO salt

**Figure 2:**
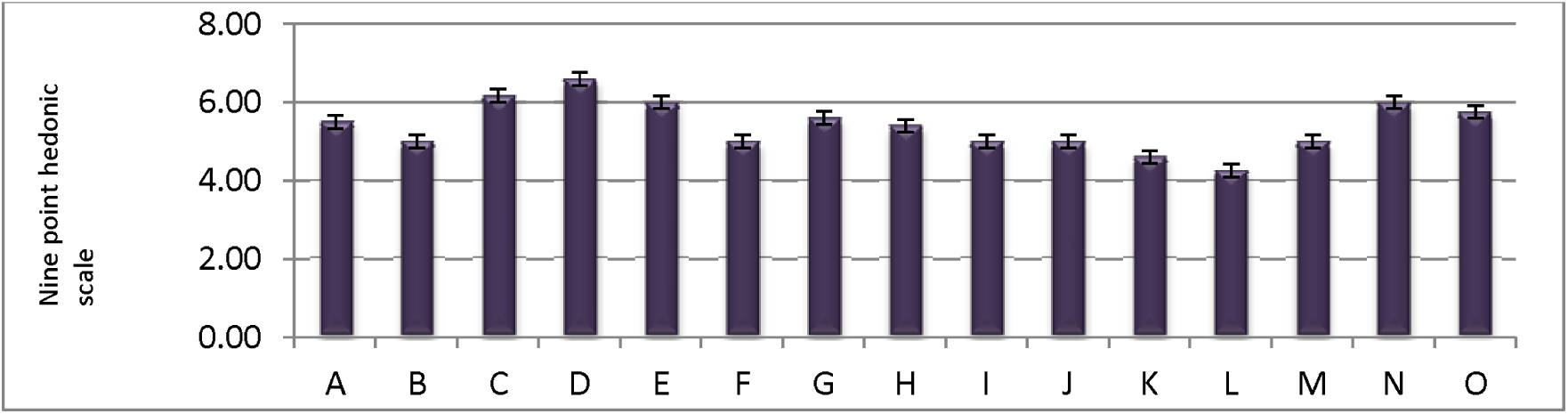
Effect of different salts on sausage tenderness. **A=** 100% Sodium chloride, **B =** 50%Sodium chloride & Potassium chloride, **C =** 50% Sodium chloride & Potassium lactate, **D=** 50% Sodium chloride & Calcium ascorbate, **E =** 25% sodium chloride, potassium chloride, Potassium lactate & Calcium ascorbate each, **F=** 50% Potassium chloride and Potassium lactate each, **G=** 50% potassium chloride &Calcium ascorbate each and **H =** 50% sodium chloride, 25% Potassium lactate & Calcium ascorbate, **I=** 25 % potassium chloride, Calcium ascorbate& 50 % Potassium lactate, J = 25 % potassium chloride, Potassium lactate & 50% Calcium ascorbate, **K =** 100% potassium chloride, **L=** 100% Potassium lactate, **M=** 100% Calcium ascorbate, **N=** 50% Potassium lactate & Calcium ascorbate, **O**= NO salt

**Figure 3:**
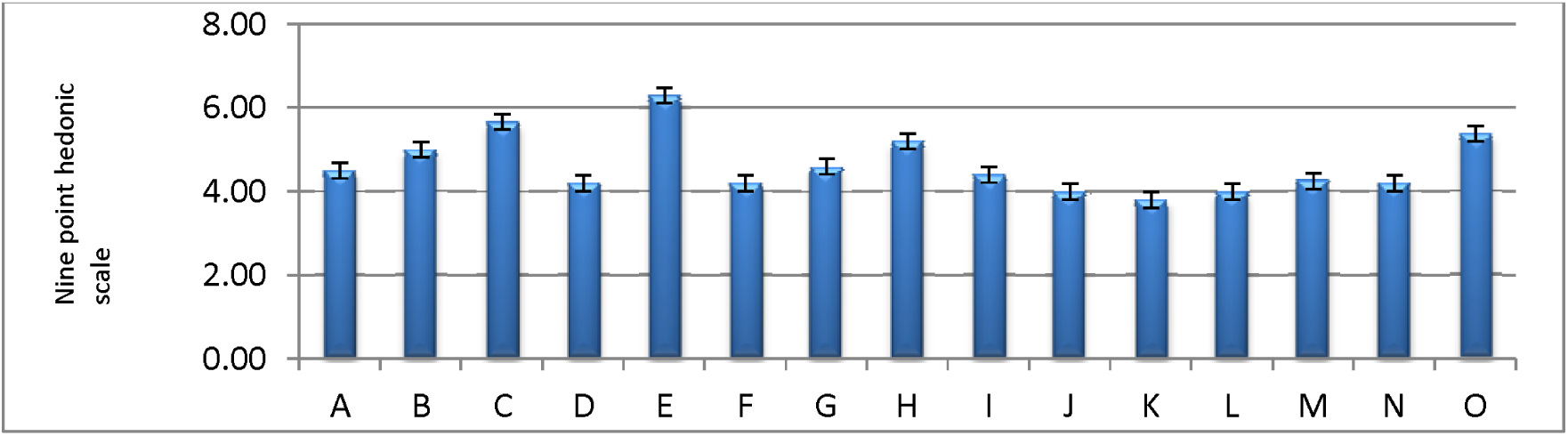
Effect of different salts on sausage juiciness. **A=** 100% Sodium chloride, **B =** 50%Sodium chloride & Potassium chloride, **C =** 50% Sodium chloride & Potassium lactate, **D=** 50% Sodium chloride & Calcium ascorbate, **E =** 25% sodium chloride, potassium chloride, Potassium lactate & Calcium ascorbate each, **F=** 50% Potassium chloride and Potassium lactate each, **G=** 50% potassium chloride &Calcium ascorbate each and **H =** 50% sodium chloride, 25% Potassium lactate & Calcium ascorbate, **I=** 25 % potassium chloride, Calcium ascorbate& 50 % Potassium lactate, J = 25 % potassium chloride, Potassium lactate & 50% Calcium ascorbate, **K =** 100% potassium chloride, **L=** 100% Potassium lactate, **M=** 100% Calcium ascorbate, **N=** 50% Potassium lactate & Calcium ascorbate, **O**= NO salt

**Figure 4:**
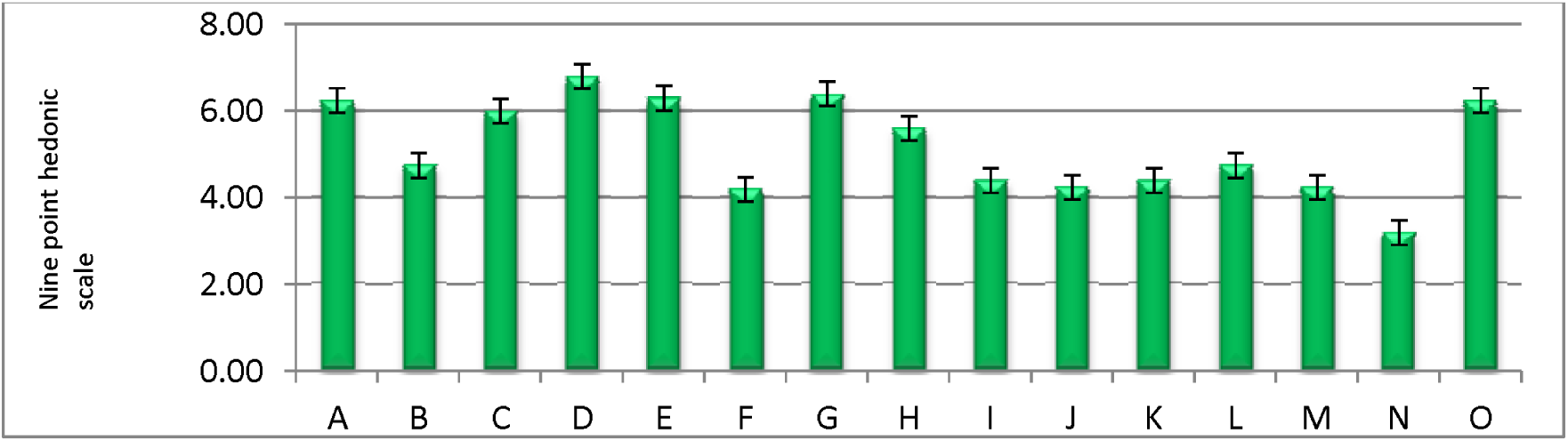
Effect of different salts on overall acceptability. **A=** 100% Sodium chloride, **B =** 50%Sodium chloride & Potassium chloride, **C =** 50% Sodium chloride & Potassium lactate, **D=** 50% Sodium chloride & Calcium ascorbate, **E =** 25% sodium chloride, potassium chloride, Potassium lactate & Calcium ascorbate each, **F=** 50% Potassium chloride and Potassium lactate each, **G=** 50% potassium chloride &Calcium ascorbate each and **H =** 50% sodium chloride, 25% Potassium lactate & Calcium ascorbate, **I=** 25 % potassium chloride, Calcium ascorbate& 50 % Potassium lactate, J = 25 % potassium chloride, Potassium lactate & 50% Calcium ascorbate, **K =** 100% potassium chloride, **L=** 100% Potassium lactate, **M=** 100% Calcium ascorbate, **N=** 50% Potassium lactate & Calcium ascorbate, **O**= NO salt.

## Discussion

Changes in pH, WHC, and rheological properties are reported to affect storage and processing quality of the meat. The microbiological stability of high pH meat is poor, tenderness is more variable, and cooked flavour is inferior (Braggins, 1996).

In relation to the acidification process the partial and total replacement of sodium chloride by potassium chloride, potassium lactate and calcium ascorbate range 5.11% - 6.77% as shown Table 4, across and among the varying inclusion level of salts and storage days.

No salt breakfast sausage had the highest pH value at day 10 while 100% calcium ascorbate had the lowest at day 0. Consistent pattern was observed as storage days increases there was increased pH value. Similar result was reported (Hand *et al*., 1982; Gelabert *et al*.,2003 and Ruusunen *et al*., 2003, while Choi *et al*.,2014 reported higher pH value ranged 6.38 – 7.80. In contrast, Gimeno *et al*. (2001b) reported a less pH value which ranged between 4.42 – 4.66. Differences in pH could be partially explained by the lactic acid bacteria development. Salt combination of 50% sodium chloride and calcium ascorbate had the highest cooking yield at day 5 compared to others, while 100% potassium lactate had the lowest value. Wettasinghe and Shahidi (1997) investigated the oxidative stability, cooking yield, and texture of low pork treated with low sodium salt mixture consisting of NaCl, KCl, MgSO4, and lysine hydrochloride. The cooking yield of products containing 1% low sodium salt mixture was lower than the product with 1% sodium chloride; however, at 2 and 3% level of mixture, it was similar to that of the product with 2 and 3% sodium chloride. The result obtained in this study could be due to the synergy of sodium chloride and calcium ascorbate to increase retention of water by the protein structure in the presence of chloride ions. The chloride ion is much more important than the sodium ion for achieving increased water binding by meat protein.

Cooking loss is a combination of liquid and soluble matter which is lost from the meat and meat product such as breakfast sausage during cooking. Cooking loss obtained in this study ranges from 11.72% - 34.86% as shown Table 6, was far below cooking loss reported by Choi *et al*. (2014) which ranges from 2.33% - 3.19%. Since salt (NaCl) aid water binding through the help of soluble protein. The high level of cooking loss obtained in this study was found in sausage without salt. This could be due to lack of salt in the product.

**Table 6:**
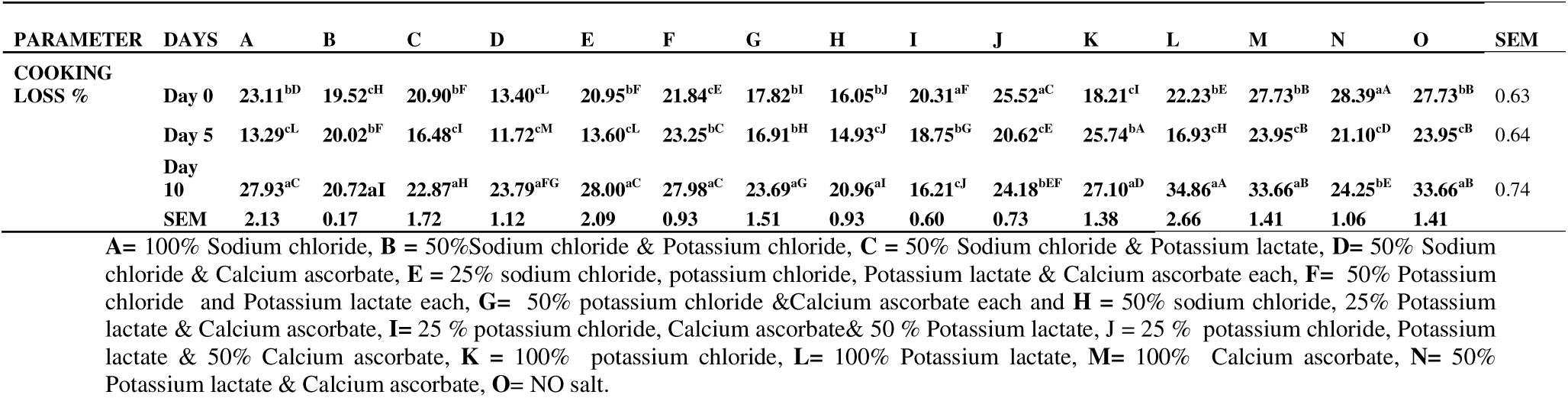
Interaction effect of different salt and storage days on cooking loss of beef sausage

The panelists were able to distinguish a difference in colour, tenderness and overall acceptability among the treatment (Figure 1 - 5). The effects of replacing NaCl by KCl, potassium lactate and glycine on physicochemical, microbiological, and sensory properties of fermented sausages were studied.. In this study, the combination of 50% sodium chloride and calcium ascorbate was rated high which could be attributed to calcium ascorbate ability to fix colour and the synergy between sodium chloride and calcium ascorbate reduced spiciness of the breakfast sausage thus making the product more tender. There were no significant differences in the microbial load. This could be due to the ability of salt to reduce or inhibit the growth of microbes (Oshibanjo, 2017).

Gelabert et al., 2003 reported that replacement of 50% of the NaCl with KCl in frankfurter provided acceptable flavour without excessive bitterness, while Hand et al., 1982a observed similar results by replacing 35% of NaCl with KCl. Hand et al., 1982b investigated the effects of chloride salts of potassium and magnesium by replacing all (100%) or part (35%) of NaCl in mechanically deboned turkey frankfurters. At substitution levels, higher than 30% with potassium lactate and higher than 50% with glycine, a loss of cohesiveness was noticed by the sensory analysis in fermented sausages.

## Conclusion

Sodium chloride replacement with Calcium Ascorbate at 50% enhanced the most preferred sausage for overall acceptability and aerobic bacteria count lower than the international standard limit.

## Acknowledgements

I acknowledged the laboratory technologists of Microbiology and Animal science Department that assisted in carried out this result in the laboratory.

## Competing interests

No competing interests declared.

## Funding

This research received no specific grant from any funding agency in the public, commercial or not-for-profit sectors’.

